# Injectisome assembly primes *Pseudomonas aeruginosa* for Type III secretion

**DOI:** 10.1101/2025.09.29.679233

**Authors:** Kristen Ramsey, Shoichi Tachiyama, Apolline Brossard, Zhao Hang, Jun Liu, Barbara I. Kazmierczak

## Abstract

Many Gram-negative pathogens, including *Pseudomonas aeruginosa*, use a Type III Secretion System (T3SS) to intoxicate eukaryotic cells. The T3SS is an important virulence factor linked to increased morbidity and mortality in infections, yet its expression slows bacterial growth and activates innate immune receptors. T3SS genes are expressed heterogeneously, with T3SS-ON cells arising from ‘primed’ bacteria that express the T3SS transcriptional activator ExsA and respond immediately to T3SS activating signals. However, the mechanistic basis for priming is not known.

ExsA is part of a complex protein-sequestration network, positively regulating its own expression as well as that of its anti-activator (ExsD), its anti-anti-activator (ExsC) and ExsC’s binding partner (ExsE). These four proteins create a bistable regulatory network. We hypothesized that transcription from a cAMP-dependent promoter upstream of ExsA could drive cells into the primed state and tested this at the single cell level. Exogenous cAMP increased the proportion of primed, ExsA-expressing cells, with whole cell cryo-electron tomography demonstrating assembly of T3SS injectisomes under these conditions. Intrastrain variation in endogenous cAMP levels correlated with strain-specific proportions of primed bacteria, while genetic manipulation of adenylate cyclases and cAMP phosphodiesterase altered primed population size. Priming occurred in planktonically cultured populations and was independent of type IV pilus assembly or retraction; however, mutation of genes in the Pil-Chp system, which regulates cAMP production, changed proportions of primed cells. This work demonstrates how endogenous and exogenous cAMP inputs into a bistable regulatory switch generate subpopulations of T3SS-primed cells poised to respond to activating signals.

**Importance:** Type III Secretion Systems (T3SS) are specialized protein secretion systems that allow bacteria to inject toxins into eukaryotic cells. T3SS are important virulence factors, but their expression carries a fitness cost: they slow bacterial growth and make bacteria vulnerable to detection by the innate immune system. Some pathogens, like *Pseudomonas aeruginosa,* balance the costs and benefits of T3SS expression by restricting T3SS expression to a subset of cells. T3SS-ON cells arise from ‘primed’ bacteria that express the transcriptional activator ExsA and respond immediately to T3SS activating signals. However, the mechanistic basis for priming is unknown.

In this study we tested whether expression of ExsA from a cAMP-dependent promoter could drive cells into the primed state and found this to be true. Whole cell cryo-electron tomography demonstrated that bacteria assembled T3SS injectisomes under these conditions. This work demonstrates how cAMP inputs into a bistable regulatory switch generate subpopulations of T3SS-primed cells.

## Introduction

Genetically identical cells, be they bacteria or cancer cells, exhibit non-identical behaviors (1, 2). Such phenotypic heterogeneity can arise from stochastic variation in gene expression that is then propagated and amplified by regulatory networks, yielding distinct subpopulations of cells (2-4). The generation of phenotypically diverse cells (e.g. antibiotic persisters) can increase adaptive fitness in unpredictable, fluctuating environments (bet-hedging) or enable a division of labor between subpopulations producing beneficial yet costly shared goods (5, 6). Among pathogenic bacteria, heterogeneous expression of the Type III Secretion System (T3SS) leads to ‘cooperative virulence’, allowing populations to balance the individual fitness cost of T3SS expression with the shared benefits of niche creation, barrier disruption and immune defense (7-9).

The T3SS injectisome is a virulence-associated nanomachine that directly injects bacterial effector proteins into host cells. This needle-like structure is composed of ∼20 proteins that are conserved among many Gram-negative pathogens and symbionts. Cryo-electron microscopy (cryo-EM) and cryo-electron tomography (cryo-ET) studies have revealed that the injectisome is composed of a cytoplasmic complex, sorting platform, envelope-spanning basal body, needle, and needle tip complex (10-14). When the needle tip complex contacts a host cell membrane, the translocon complex is recruited at the tip, forming a channel into the host cell (15). Translocated effectors then target a variety of host cell functions which promote bacterial survival, inhibit immune activation and often result in host cell death. In the opportunistic pathogen *Pseudomonas aeruginosa*, the injectisome targets eukaryotic cells ranging from predatory amoeba to human phagocytes and epithelial cells, leading to cytoskeletal disruption and cell death (16-20). The T3SS injectisome is critical for *P. aeruginosa* to establish acute murine and human infection and is linked with increased morbidity and mortality in hospital-acquired infections (21, 22).

Single-cell studies have established that *P. aeruginosa* heterogeneously expresses T3SS genes, yielding bimodal populations of “OFF” and “ON” cells in the presence of activating signals (23-27). T3SS-ON cells grow slowly, while recognition of T3SS components by NLRC4 inflammasomes may incur additional fitness costs during infection (26, 28, 29). Nonetheless, ExoU, a phospholipase A2 injected by the T3SS, rapidly kills recruited neutrophils and macrophages and serves as a shared public good in murine infections (20, 23). Thus *P. aeruginosa* may use phenotypic heterogeneity to maintain the T3SS+ genotype while avoiding the measurable fitness costs associated with expressing this virulence trait (23, 26).

The AraC-family transcriptional activator, ExsA, controls the entire *P. aeruginosa* T3SS regulon, which encodes the needle complex, secreted effectors, and ExsA regulators (30, 31). Upregulation of ExsA-dependent gene expression is coupled to activation of the T3SS injectisome via a “partner-switching” model that assumes the presence of an assembled T3SS needle. In the absence of activating signals, the needle is closed and incapable of secretion; intracellular ExsA is bound by its anti-activator ExsD, and ExsC is sequestered by ExsE (32, 33). Activation signals, such as host cell contact or *in vitro* Ca^+2^ chelation, induce translocon formation and allow ExsE secretion, freeing ExsC to bind ExsD (33-35). This allows ExsA to dimerize and promote transcription of the T3SS regulon. Such a model raises the question, however, of how T3SS genes are expressed to allow assembly of the injectisome in the first place.

Prior work has established that, naïve of any exogenous T3SS-activating signal, a subpopulation of cells expresses high levels of ExsA (26). When exposed to a T3SS-activating signal, these ExsA-positive cells rapidly gave rise to most T3SS-expressing cells in the population, behaving as if they were ‘primed’ to respond. In this study we examined how priming arises and whether it is correlated with T3SS needle assembly. We also investigated the role of the second messenger cyclic-AMP (cAMP), which activates a Vfr-cAMP dependent promoter upstream of *exsA* (P*_exsA_*) that drives ExsA expression but not that of ExsD, ExsC or ExsE (25). cAMP is a well-established positive regulator of *P. aeruginosa* T3SS gene expression: deletion of genes encoding *vfr*, the adenylate cyclases *cyaA* and/or *cyaB*, or P*_exsA_* itself significantly impair T3SS gene expression and reduce *P. aeruginosa* virulence (25, 36, 37). Given the importance of cAMP to T3SS-dependent virulence, we chose to test its role in generating heterogeneous T3SS priming and expression.

## Results

### cAMP is sufficient to increase the proportion of primed cells

Lineage tracing of *P. aeruginosa* PA14 cells carrying chromosomal reporters for ExsA production (*exsA*-IRES-mTagRFP-t) and ExsA-dependent T3SS gene transcription (*attB*::P*_exoT_*-sfGFP) has demonstrated that an ExsA+/RFP+ subpopulation of cells is present even in the absence of T3SS-activating signals (e.g. in Ca^+2^ replete conditions), and that these cells quickly respond to activating signals (such as the chelator nitriloacetic acid (NTA)) by inducing T3SS gene expression (26). We used this dual reporter strain to test whether cAMP alters priming and/or T3SS gene expression, measuring individual cell fluorescence by flow cytometry. In MinS + Ca^+2^ media, treatment with cAMP increased ExsA-expressing primed cells from 3.5% to ∼70% of the population (Figure 1A). Expression of the P*_exoT_* transcriptional reporter remained low in the absence of T3SS-activating signals regardless of cAMP exposure (Figure 1A). In T3SS-activating MinS, cAMP treatment also increased the percentage of ExsA+/RFP+ cells. As expected, P*_exoT_* expression increased in tandem (Figure 1B). Thus, cAMP treatment was sufficient to increase the proportion of ExsA-expressing cells, but an activating signal was still required for upregulation of T3SS expression.

**Figure 1.**
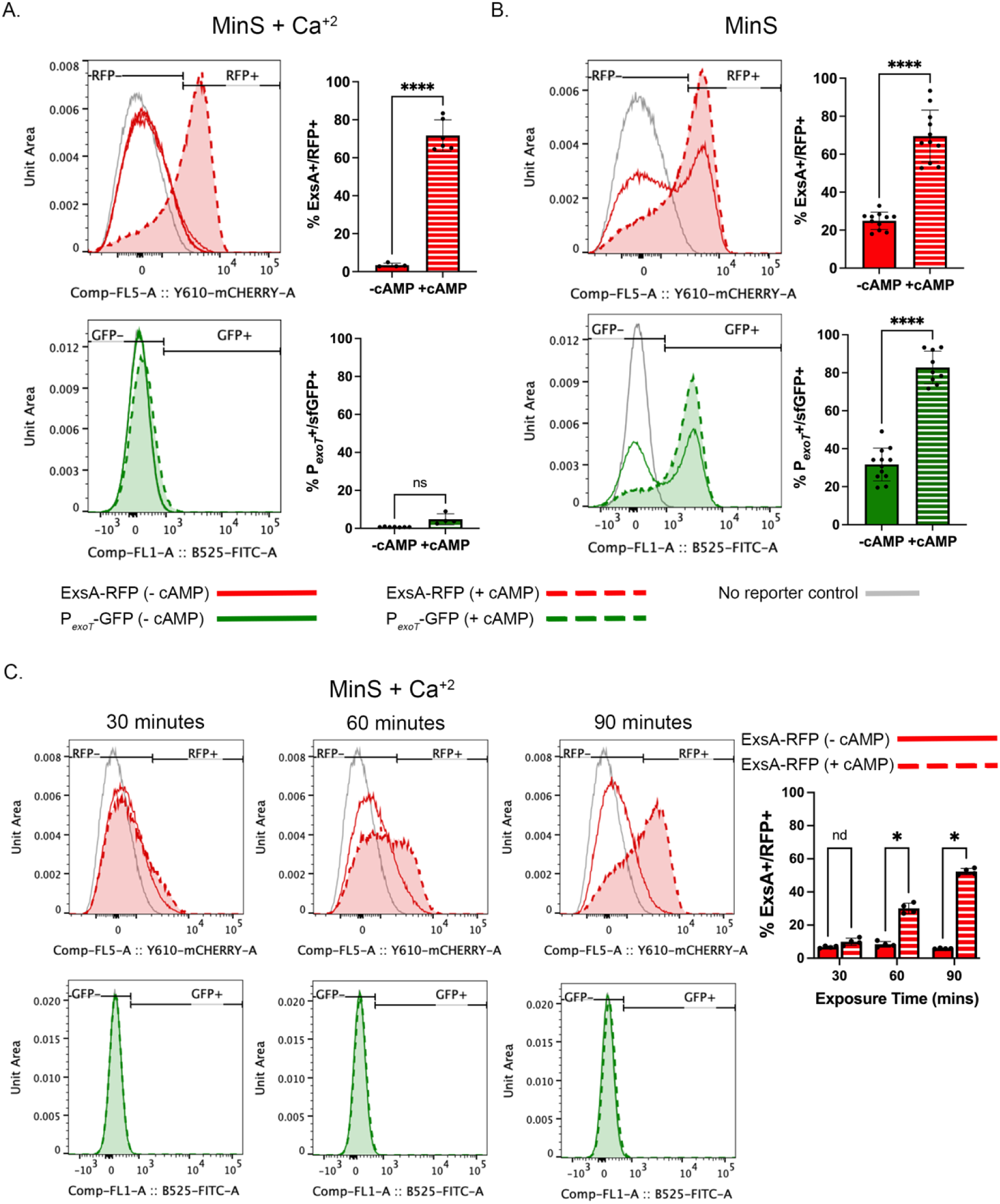
Exogenous cAMP is sufficient to increase ExsA+, T3SS-primed PA14 cells. Representative flow cytometry histograms measuring ExsA production (red, RFP+) and ExoT transcription (green, GFP+) in PA14 dual ExsA/T3SS reporter cells grown planktonically for 7 hours in (A) T3SS secretion-repressing MinS + 5 mM Ca^+2^ media or (B) T3SS secretion-activating MinS media with addition of vehicle (solid) or 20 mM cAMP (dashed), quantified over all experiments (n=4-12) (right). Significance determined by Welch’s unequal variances *t*-test. (C) Representative flow cytometry histograms measuring ExsA production (RFP) and ExoT transcription (GFP) of PA14 dual ExsA/T3SS reporter cells grown in MinS + Ca^+2^ pulsed at t=2 hours with vehicle (solid) or cAMP (dashed) for either 30-, 60-, or 90-minutes during outgrowth. Cultures were examined by flow cytometry at 7 hours of total MinS + Ca^+2^ growth. Significance determined by Welch’s unequal variances *t*-test (n = 3-4). Fluorescent gates were established based on a *P. aeruginosa* PA14 control with no fluorescent reporter (grey). Bars show mean ± S.D. P-values: *, <0.05; **** <0.0001.

Intracellular [cAMP] increases in *P. aeruginosa* cells upon surface contact via the attachment/retraction of Type 4 pili (T4P) and the activation of the Pil/Chp chemotaxis system (38, 39). Individual cells appear to have a multigenerational memory of intermittent surface attachment, leading to variation in cAMP reporter expression among cells over time (39-41). We wondered whether a brief exposure to cAMP would be sufficient to increase the ExsA+ subpopulation several hours later. Indeed, exposing PA14 dual reporter cells to cAMP for as little as 30 minutes increased the percentage of ExsA+ cells in MinS + Ca^+2^, as compared to vehicle control (Figure 1C). Thus a brief pulse of cAMP could increase numbers of ExsA+/RFP+ cells, again without resulting in strong induction of the P*_exoT_*-sfGFP reporter under these secretion-repressing conditions.

### The T3SS needle is assembled in the absence of activating signals

The Vfr-cAMP dependent P*_exsA_* promoter is uniquely capable of expressing *exsA* in the absence of any of ExsA’s partner-switching regulators. We wondered whether this generated enough free ExsA to drive expression and assembly of the T3SS, implying that a subpopulation of cells would assemble T3SS needles under T3SS repressing conditions. To test this, we visualized surface structures of live *P. aeruginosa* PA14 and PA103 with whole-cell cryo-ET. We could readily identify and count needles (Figure S1); subtomogram averaging on these needle-like complexes allowed us to determine *in situ* structures and confirm visualization of injectisomes with basal bodies in the periplasm (Figure S1). When grown in MinS+Ca^+2^, five of 36 (13.8%) PA14 cells and over 75% of PA103 cells had one or more T3SS needles (Figure 2). PA103, which has a high proportion of T3SS-ON cells (26) also had more needles per cell (mean, 3.5 ±1.9) than PA14. As expected, T3SS effector secretion was repressed under these conditions where needle assembly was observed (Figure S2). Similar proportions of cells assembled injectisomes when PA14 (16.7%) and PA103 (76.9%) were cultured in T3SS-activating MinS+ NTA media (Figure 2).

**Figure 2.**
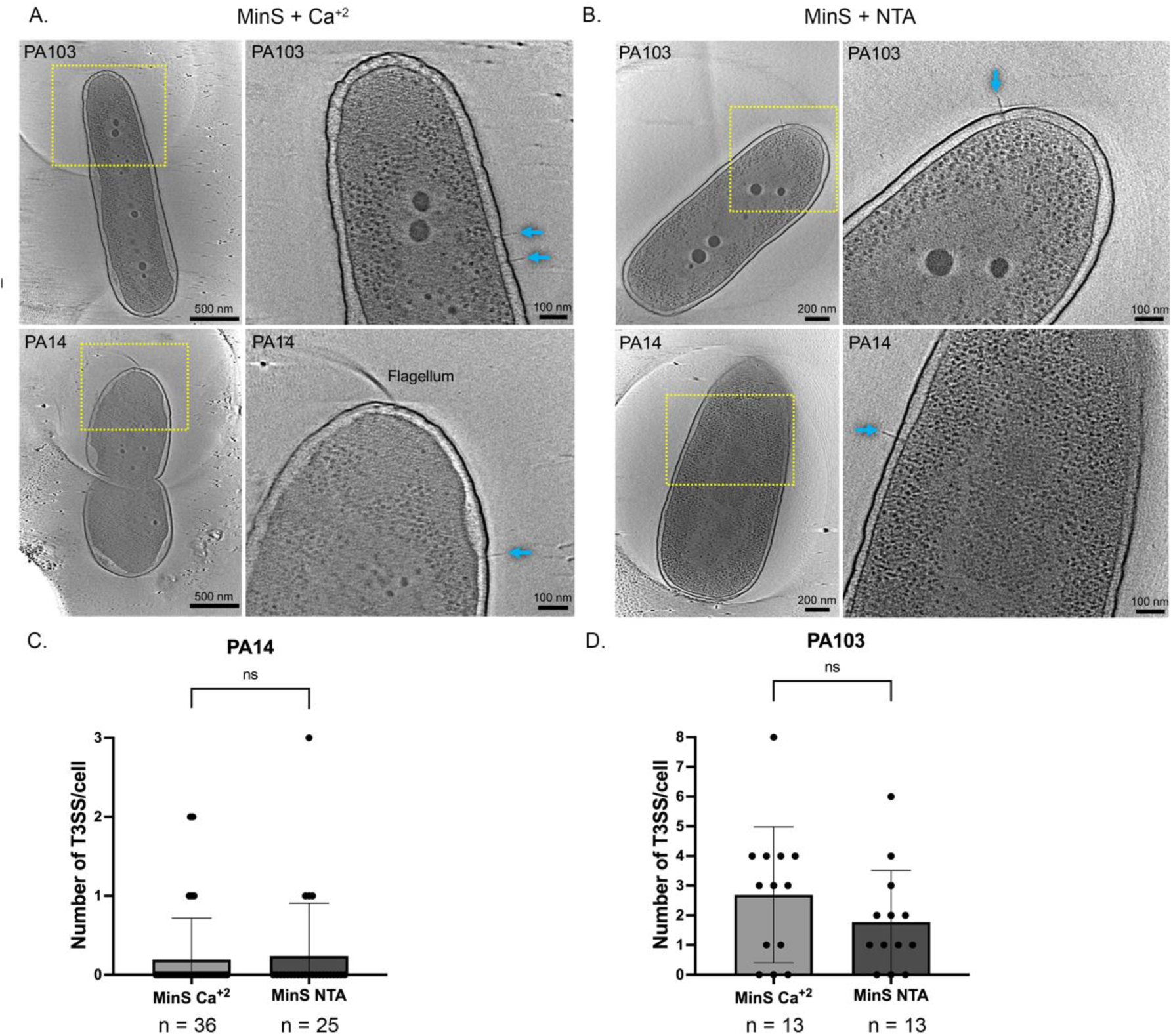
T3SS injectisome assembly is not dependent on an activation signal. PA103 and PA14 WT cells were grown in (A) MinS + 5 mM CaCl_2_ or (B) MinS + 10 mM NTA for 7 hours, then prepared and imaged by cryo-ET at a magnification of 6.184 Å physical pixel size. Type III injectisomes on the bacterial surface were counted manually and are indicated by blue arrows. Representative tomograms shown. (C) Quantification of PA14 T3SS needles per cell in both growth conditions (n = 25-36). (D) Quantification of PA103 T3SS needles per cell in both growth conditions (n = 13). Significance determined by Welch’s unequal variances *t*-test; ns, not significant.

Treatment with cAMP greatly increased the proportion of ExsA+/RFP+ ‘primed’ cells as measured by flow cytometry; we examined whether this would also increase the number of cells with assembled T3SS needles. Whole-cell cryo-ET demonstrated that 93% of PA14 cells cultured in MinS+Ca^+2^ plus cAMP assembled T3SS needles, with an average of 5.2 needles per cell (Figure 3A-B). Thus a subset of bacteria assembled T3SS needles in the absence of T3SS activating signals, and cAMP-induced expression of ExsA allowed for a “burst” of T3SS gene expression sufficient for T3SS needle assembly.

**Figure 3.**
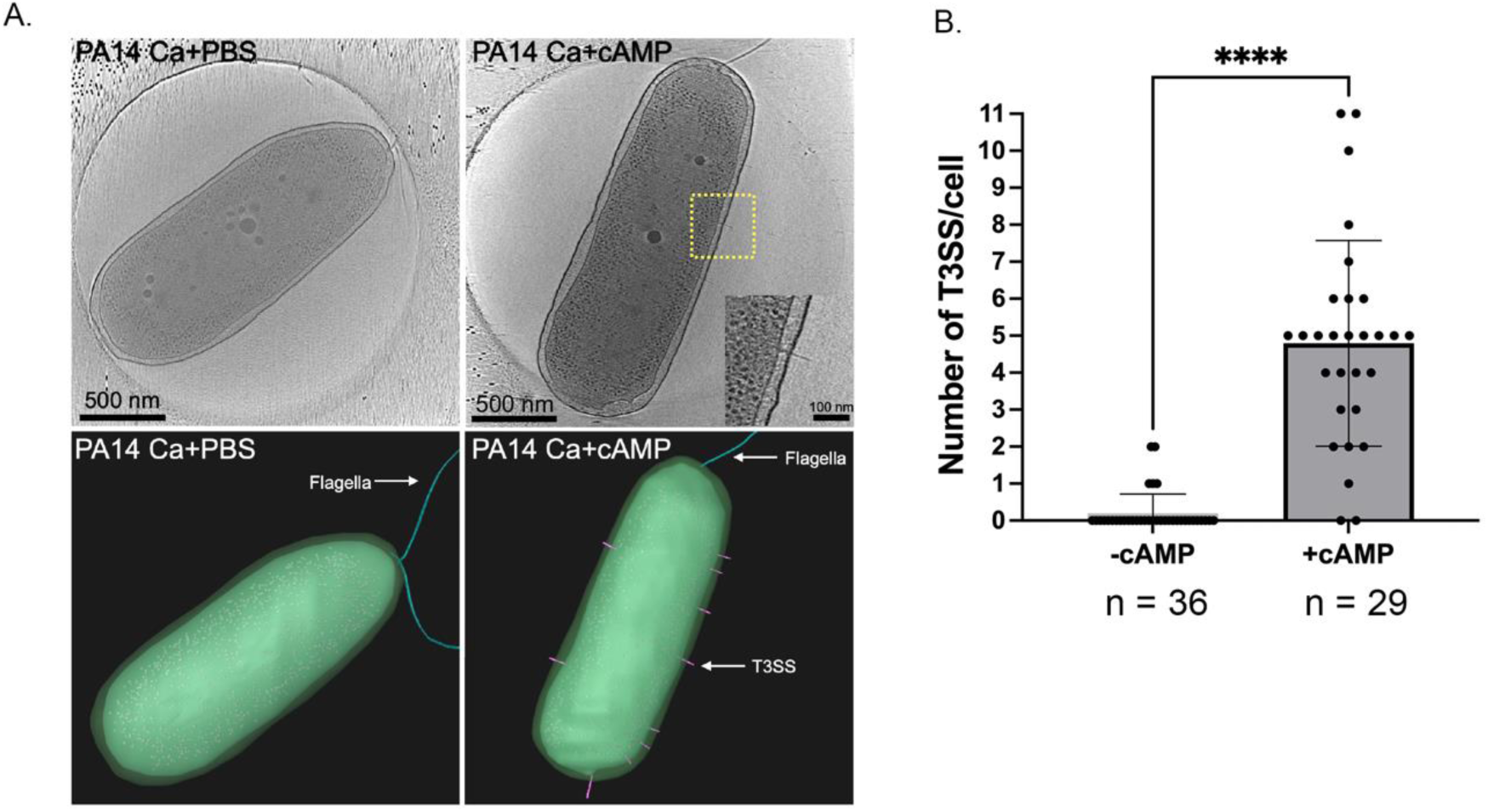
Exogenous cAMP increases proportion of PA14 cells assembling injectisomes in MinS + Ca^+2^. (A) Example tomogram of a PA14 cell grown in MinS + Ca^+2^ in the absence (left) or presence (right) of 20 mM cAMP for 7 hours, then prepared and imaged by cryo-ET at a magnification of 6.184 Å physical pixel size. Inset: magnification of a representative injectisome in additional structural detail. 3D composite tomograms of each cell are shown below with the T3SS injectisome pseudocolored pink. (B) Quantification of T3SS injectisomes per cell grown in MinS + Ca^+2^ in the absence (left) or presence (right) of cAMP. Significance determined by Welch’s unequal variances *t*-test (n = 29-35). Bars represent the mean ± S.D. of all individual points. **** = P-value <0.0001.

### Interstrain variation in endogenous cAMP reflects differences in T3SS-ON populations

Most PA103 cells grown in the absence of cAMP treatment assembled T3SS needles, often several per cell. We also observed many T4P on these cells by cryo-ET, even though cells were cultured planktonically, when T4P are usually not assembled (42) (Figure S3). Increased assembly of a second nanomachine positively regulated by cAMP suggested that endogenous cAMP levels were higher in PA103 relative to PA14; this was tested at the single-cell level by integrating a cAMP-responsive *lacP1*-sfGFP transcriptional reporter into each background at the chromosomal *attB* site (43). This reporter responded robustly to exogenous cAMP (Figure S4), as previously observed (38). *lacP1*-reporter fluorescence distributions were rightward skewed in both PA103 and PA14 (Figure 4A-B), with median fluorescence intensity (2185.5) of PA103 significantly greater than that of PA14 (180.45, *p* = 0.014). Under these T3SS activating conditions (MinS + NTA), almost all PA103 cells expressed the P*_exoT_* reporter, while PA14 P*_exoT_* reporter expression was bimodal (Figure 4A-B). Measurements of cAMP and P*_exoT_*expression were also extended to two additional *P. aeruginosa* strains, PAK and PAO1. Unlike PA103 and PA14, these well-characterized isolates do not secrete the T3SS effector ExoU, instead translocating ExoS along with ExoT and ExoY into target cells (44-46). *lacP1* reporter fluorescence was bimodally distributed in planktonically grown PAK cells; the median fluorescence intensity of the population (1628) was significantly higher than that observed for PAO1 (152.9, p = 0.027) (Figure 4C-D, S5). Both PAO1 and PAK showed bimodal expression of P*_exoT_*-GFP when planktonically grown (Figure 4C-D). Expression of *lacP1*-sfGFP was low in the adenylate cyclase mutant PAK Δ*cyaA*Δ*cyaB*, as expected. Thus cAMP levels were overall higher in strains with a higher proportion of P*_exoT_*-GFP+, T3SS-ON cells (i.e. PA103 and PAK). However, the intensity and distribution of *lacP1*-GFP fluorescence was not linearly reflected in P*_exoT_* reporter activity. This is consistent with cAMP serving as a signal that “flips the switch” to push cells into a bistable ExsA-expressing, ‘primed’ state, rather than as a signal that drives expression of T3SS genes *per se* (47).

**Figure 4.**
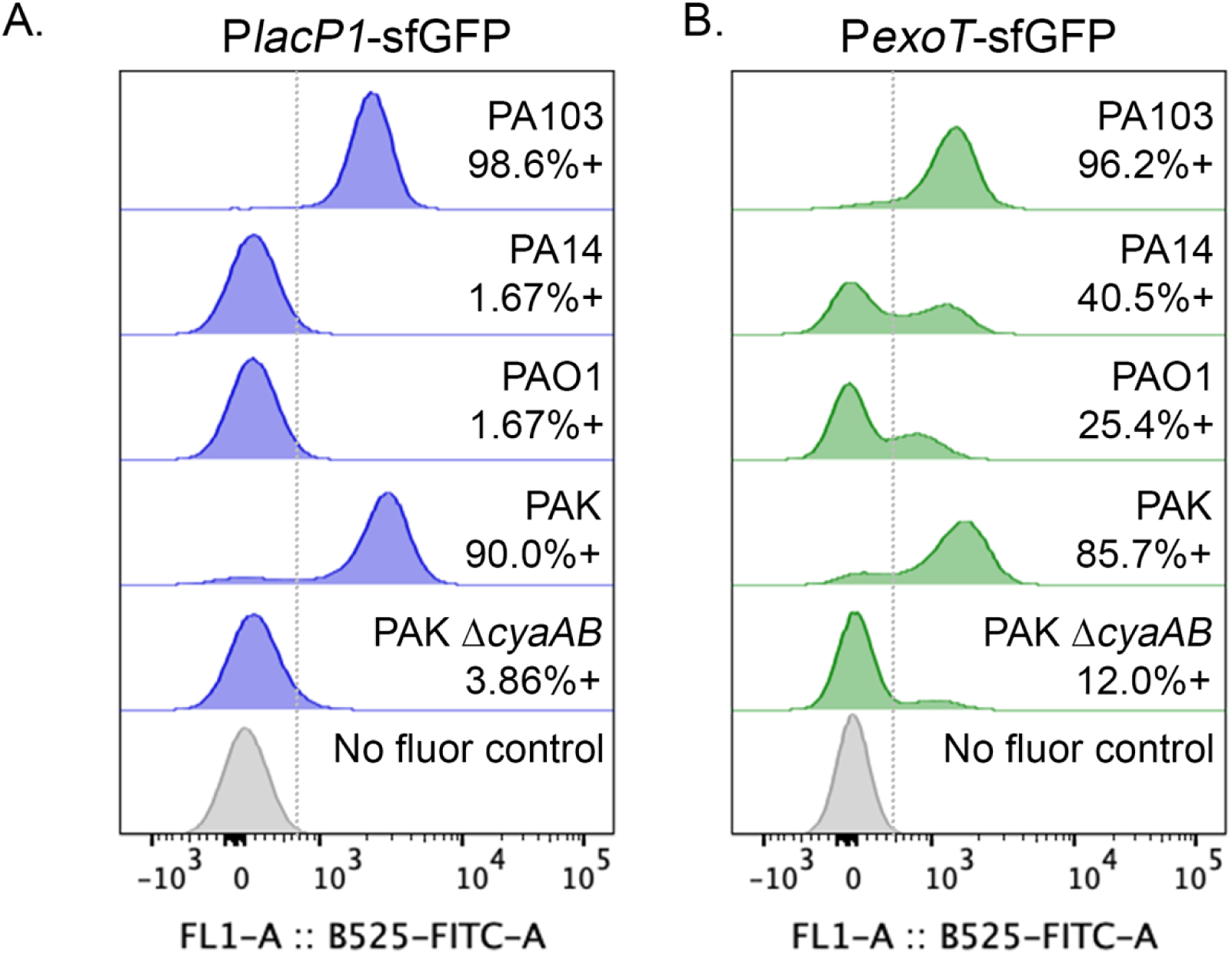
Variation in endogenous cAMP levels correlates with the proportion of T3SS+ cells. PA103, PA14, PAO1, PAK, and PAK Δ*cyaAB* carrying either the cAMP responsive *attB*::*lacP1*-sfGFP transcriptional reporter (A) or the *attB*::P*_exoT_*-sfGFP T3SS transcriptional reporter (B) were cultured planktonically for 7 hours in MinS + NTA, fixed, and analyzed for GFP fluorescence by flow cytometry, with fluorescent gates established based on a *P. aeruginosa* PA14 control with no fluorescent reporter (grey histogram/line).

### Mutations that modulate endogenous cAMP levels alter the proportion of T3SS-ON cells

*P. aeruginosa* cAMP levels are controlled by two adenylate cyclases, CyaA and CyaB, and the phosphodiesterase CpdA (36, 48). We tested whether the high proportion of T3SS-ON PA103 bacteria could be reduced by overproduction of the CpdA phosphodiesterase, and found this to be the case (Figure 5). Likewise, the frequency of ExsA+ and T3SS-ON PA14 bacteria were reduced by overproduction of the CpdA phosphodiesterase (Figure 5).

**Figure 5.**
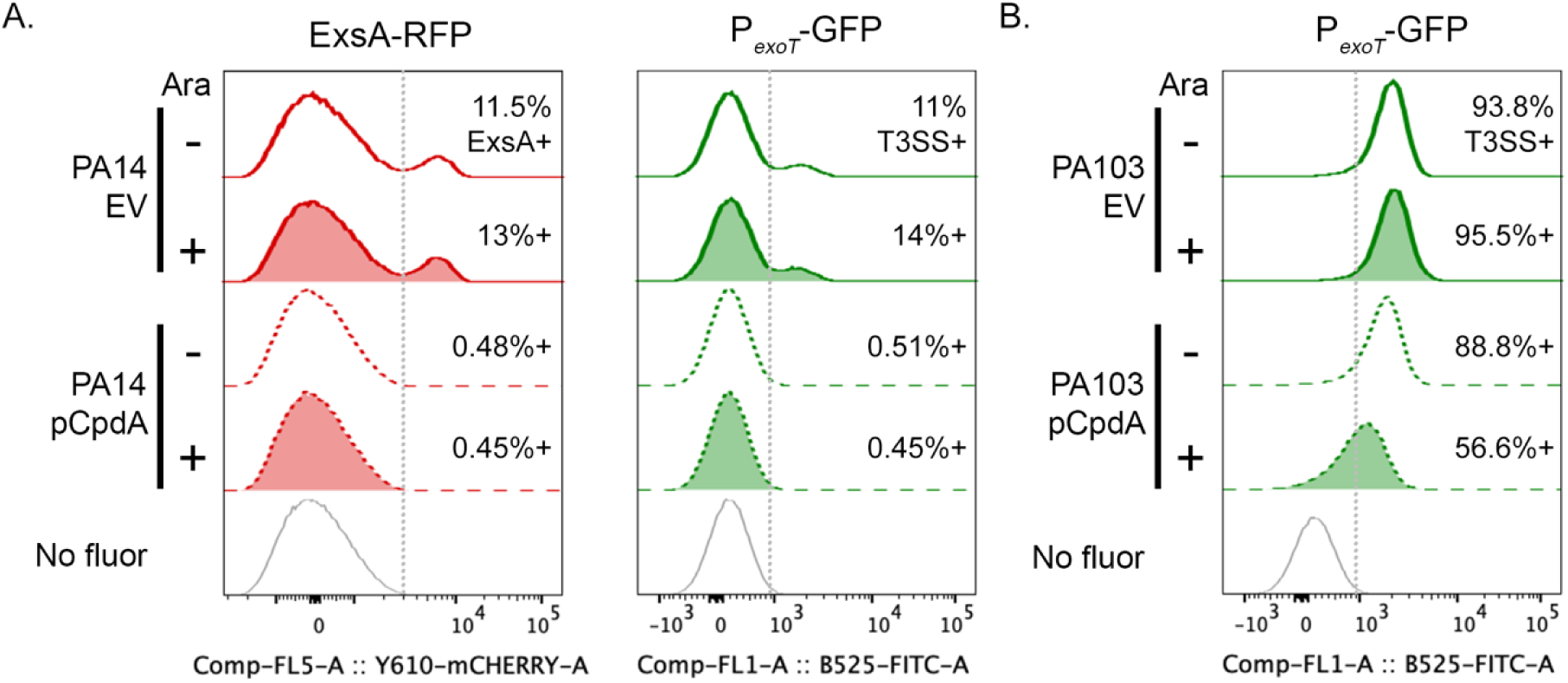
Breakdown of cAMP decreases T3SS-priming. (A) pMQ72 (“EV”, solid line) or pCpdA (dotted line) was introduced into the PA14 *exsA*-IRES-mTagRFP-t P*_exoT_*-sfGFP T3SS dual reporter strain. Bacteria were cultured planktonically in MinS for 7h, then analyzed for RFP (left, red) and GFP fluorescence (right, green) by flow cytometry, with fluorescent gates established based on a *P. aeruginosa* PA14 control with no fluorescent reporter (grey line). Expression of CpdA was induced by adding 0.4% arabinose to MinS as indicated (+, filled histograms). (B) PA103 *attB*:: P*_exoT_*-sfGFP T3SS reporter plus pMQ72 (“EV”, solid line) or pCpdA (dotted line) was grown and analyzed as in (A).

Although CyaA and CyaB each contribute measurably to cAMP production, CyaB is the primary target of the Pil/Chp chemosensory pathway, producing cAMP in response to T4P extension, binding and retraction (Figure S6) (38, 39, 49). As cryo-ET imaging showed assembled T4P even when PA103 was cultured planktonically, we used the P*_exoT_* reporter to determine whether disruption of T4P assembly proteins would decrease the proportion of T3SS-ON bacteria in liquid grown populations (50). We observed no change in the proportion of P*_exoT_*-GFP+ cells between wild-type PA103, Δ*pilQ* (lacking the T4P secretin), Δ*fimX* (deficient in T4P assembly) or Δ*pilT* (lacking the primary retraction ATPase) (Figure S7). A similar pattern was seen when this reporter was introduced into a series of PA14 Tn insertion mutants targeting proteins required for T4P assembly or retraction (Figure 6A). In contrast, disruption of the major pilin subunit (Δ*pilA*) or the chemoreceptor protein (Δ*pilJ*) significantly decreased the proportion of T3SS-ON cells (Figure 6B-C).

**Figure 6.**
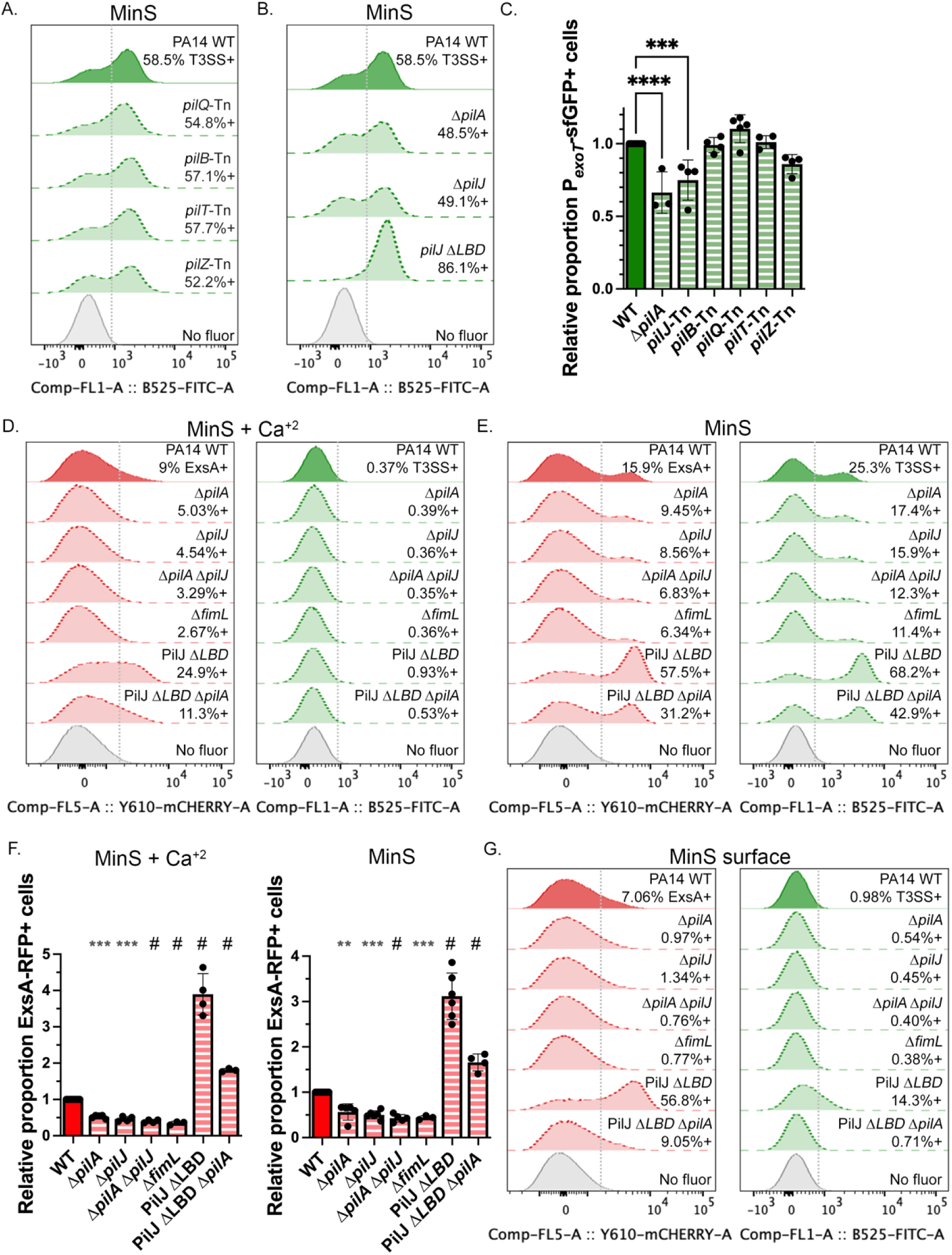
Mutation of Pil/Chp genes but not T4P assembly/retraction genes affects T3SS-priming. Representative flow cytometry histograms of PA14 WT and transposon (A) or unmarked deletion (B) T4P mutants carrying the *attB*::P*_exoT_*-sfGFP T3SS transcriptional reporter. Cells were grown planktonically for 7 hours in MinS, fixed, and analyzed for GFP fluorescence by flow cytometry. Fluorescent gates were established based on a *P. aeruginosa* PA14 with no fluorescent reporter (grey line). (C) Each point shows the proportion of T3SS+ cells, normalized to the positive WT control in each experiment, for the indicated T4P mutants; bars show mean ± S.D. of 3-5 independent experiments. Only Δ*pilA* and *pilJ*::Tn differed significantly from wild-type. (D-E) Representative flow cytometry histograms of PA14 ExsA-IRES-mTagRFP-t *attB*::P*_exoT_*-sfGFP (“WT”, solid line) and isogenic mutant strains (dotted lines) grown for 7 hours planktonically in (D) MinS + 5 mM CaCl_2_ or (E) MinS, then analyzed for RFP (left, red) and GFP fluorescence (right, green) as in (A). (F) Each point shows the proportion of ExsA/RFP+ cells in Pil/Chp mutant populations grown in MinS + Ca^+2^ or MinS, normalized to the WT control; bars show mean ± S.D. of 2-5 experiments. (G) PA14 dual reporter bacteria (“WT”, solid line) and isogenic T4P mutants (dotted lines) were harvested from MinS agar plates and analyzed for ExsA expression by flow cytometry as in (E). In panels C and F, significance was determined by one-way ANOVA with Dunnett’s multiple comparisons test comparing to WT. Adjusted p-values: * <0.05, ***= 0.0001, **** / # < 0.0001.

The PilJ chemoreceptor signal and/or ligand(s) have largely been studied in surface-associated bacteria, where the signal that upregulates cAMP production continues to be debated. T4P retraction by PilT, PilA interaction with PilJ, and ligand sensing by PilJ have all been identified as cues for pathway activation, with FimL serving as a required polar scaffold (39, 49, 51-54). We introduced unmarked deletions of *pilA*, *pilJ, fimL*, and *pilA pilJ* into the PA14 dual reporter strain and tested whether they were required for T3SS priming. Disruption of any of these Pil/Chp-associated genes reduced the proportion of ExsA+/RFP+ bacteria in the absence of a T3SS-activating signal (MinS + Ca^+2^) as well as in T3SS-activating conditions (MinS) in planktonically grown cells (Figure 6D-F). This could be rescued by providing exogenous cAMP or by complementing gene deletions *in trans* (Figure S8-S9). We also enumerated ExsA+/RFP+ cells in agar-grown WT and T4P mutant populations and observed that the loss of *pilA*, *pilJ* or *fimL* decreased the proportion of primed cells, relative to wild-type, to a greater extent than seen in planktonically grown populations (Figure 6G).

The role of PilJ’s periplasmic ligand binding domain (LBD) in cAMP signaling is still poorly understood. We deleted the region described by Yarrington *et al.* in chromosomally encoded PilJ (ΔLBD1-2, aa 39-303) in the PA14 dual reporter strain and found that this mutation increased the proportion of primed cells in all tested growth conditions, including on a surface (Figure 6B-G) (52). An *attB::lacP1*-sfGFP reporter in PA14 PilJΔLBD1-2 showed a modest but reproducible increase in cAMP levels as compared to the isogenic wild-type strain (Figure S10). Although deletion of *pilA* in the PilJΔLBD1-2 dual reporter background decreased the proportion of primed cells relative to PilJΔLBD1-2, this double mutant still had more primed cells than WT PA14. We also deleted the predicted methylesterase (ChpB) and methyltransferase (PilK) of PilJ in the PA14 dual reporter strain; their disruption respectively increases or decreases measured cAMP levels in surface-grown bacteria (55). Deletion of *chpB* increased the proportion of primed cells when cultured in either MinS + Ca^+2^ and MinS, similar to the PilJ*Δ*LBD1-2 mutant (Figure 7). A Δ*chpB ΔpilA* double mutant had fewer primed cells as compared to Δ*chpB*, but maintained increased priming relative to the wild-type dual reporter strain (Figure 7). Therefore, mutations that increase PilJ activity – and thereby adenylate cyclase activity and [cAMP] - also increased the proportion of *P. aeruginosa* primed for T3SS gene expression, and PilA affected but was not strictly required for this phenotype.

**Figure 7.**
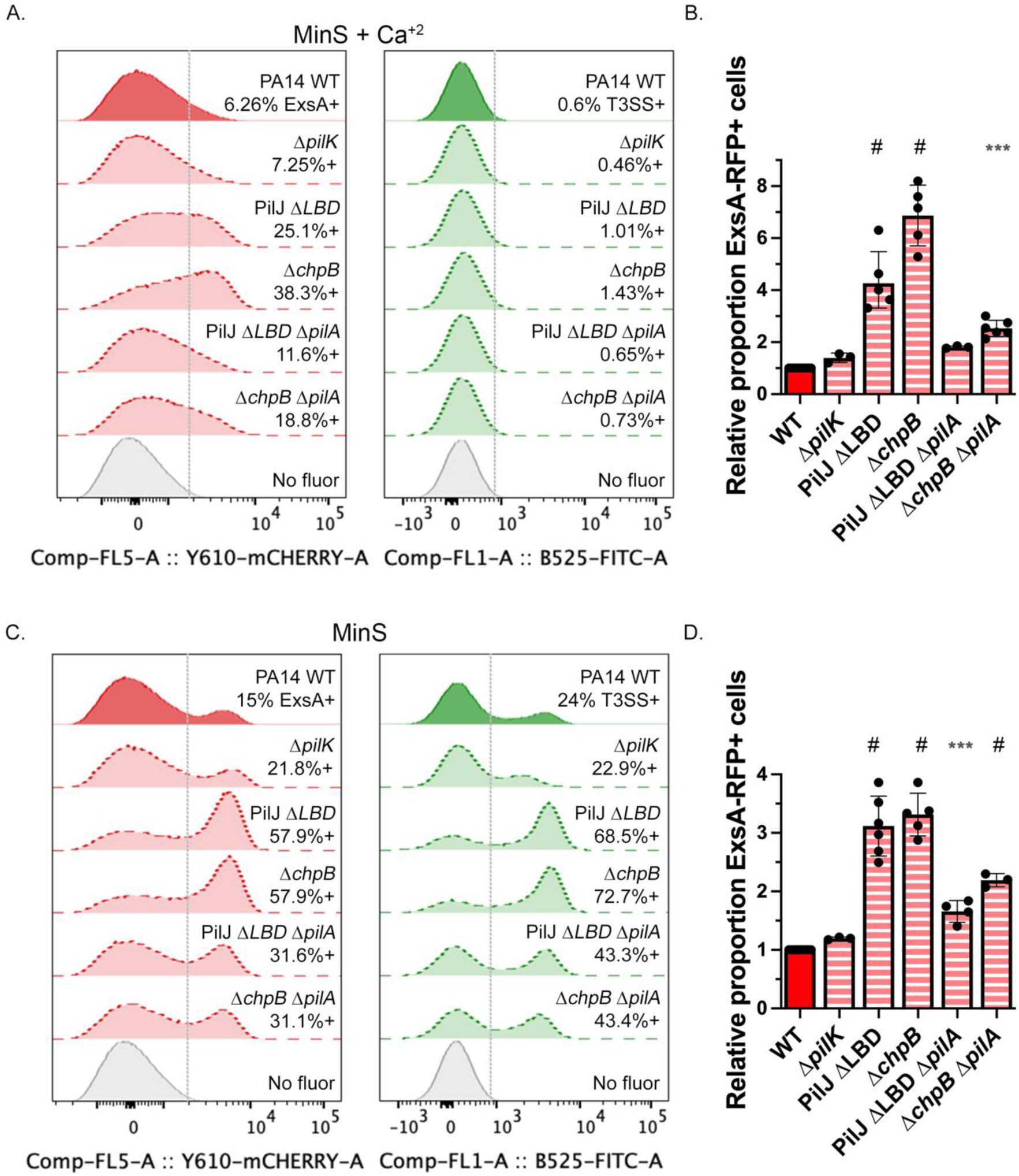
PilJ methylation state affects T3SS priming. Representative flow cytometry histograms of PA14 WT (solid line) and mutant (striped lines) bacteria carrying dual ExsA-IRES-mTagRFP-t and P*_exoT_*-sfGFP T3SS reporters grown planktonically for 7 hours in (A) MinS + 5 mM CaCl_2_ or (C) MinS, then analyzed for RFP (left, red) and GFP fluorescence (right, green) by flow cytometry. Fluorescent gates were established based on a *P. aeruginosa* PA14 with no fluorescent reporter (grey line). (B, D) Each point shows the proportion of ExsA+ cells in each population grown in (B) MinS + Ca^+2^ or (D) MinS, normalized to the WT control; bars show mean ± S.D. of 3-4 experiments. Significance determined by one-way ANOVA with Dunnett’s multiple comparisons test comparing to WT. P-values: ***, <0.001; #, < 0.0001.

## Discussion

The heterogeneous expression of Type III secretion genes within clonal populations is broadly observed. Such cooperative virulence – in which T3SS-ON individuals incur a measurable fitness cost to produce a virulence factor with shared benefit – allows genetically identical T3SS-OFF individuals to survive within a host without loss of the T3SS-positive genotype (7, 23, 56). In *P. aeruginosa*, T3SS-ON cells arise from a subpopulation of primed cells expressing the transcriptional activator ExsA. When exposed to signals that activate Type III secretion, e.g. divalent chelators such as NTA, primed cells immediately respond by expressing ExsA-dependent genes. ExsE secretion through the T3SS needle links the activation of substrate secretion with the induction of ExsA-dependent gene expression, triggering a partner-switching cascade that sequesters the anti-activator ExsD and frees ExsA to homodimerize and bind DNA (33-35). This model predicts that primed cells carry T3SS injectisomes, which we have demonstrated in this study using cryo-ET. The proportion of cells bearing injectisomes differs greatly between a strain of *P. aeruginosa* in which relatively few cells are ‘primed’ (PA14) and one in which nearly all cells exhibit a primed phenotype (PA103); PA103 cells also express more injectisomes per cell (Figure 2). As we assume that these injectisomes can be inherited from mother to daughter, having more injectisomes per cell increases the likelihood that both PA103 daughters retain this needle-positive, primed phenotype.

ExsA is encoded within the *exsCEBA* operon, and thus subject to positive transcriptional feedback; the anti-activator, ExsD, is likewise encoded within an ExsA-activated *exsDpscBCDEFGHIJKL* operon which also contains many of the T3SS injectisome structural genes. ExsA expression is therefore coupled to the expression of its anti-activator. However, Yahr and colleagues reported the presence of a weak promoter upstream of *exsA* which is activated by cAMP-bound Vfr and transcribes *exsA* alone (25). We have demonstrated that provision of exogenous cAMP is sufficient to increase the proportion of cells that express ExsA – and also to increase cells expressing the T3SS injectisome, and those that respond to a T3SS-secretion activating signal with P*_exoT_*-GFP expression. This strongly argues that primed cells carry the T3SS injectisome. We also attempted to label injectisomes on live bacteria with anti-PcrV monoclonal antibodies (as described in (27)) and then image them for P*_exoT_*-sfGFP expression by time-lapse microscopy. These experiments were unsuccessful, as our panel of monoclonal antibodies stained unfixed WT and Δ*exsA* bacteria similarly, likely a consequence of our selecting these reagents based on their performance in ELISA assays against immobilized PcrV (57).

The sequestration-based ExsA/D/C/E regulatory circuit provides an ultrasensitive mechanism for generating bistability, as has been demonstrated by reconstituting this signaling cascade in *E. coli* (58). Increased expression of free ExsA from the cAMP-Vfr dependent P*_exsA_* promoter can drive a cell across the threshold between stable OFF and ON states, resulting in the observed increases in T3SS needle-expressing, primed cells which correspond to the ON state of this bistable system (47). As illustrated in our model (Figure 8), transcriptional noise at the P*_exsA_* promoter may also lead to ExsA expression, but endogenous or exogenous cAMP signals increase the likelihood that the ExsA positive-feedback loop is triggered and the primed state established.

**Figure 8.**
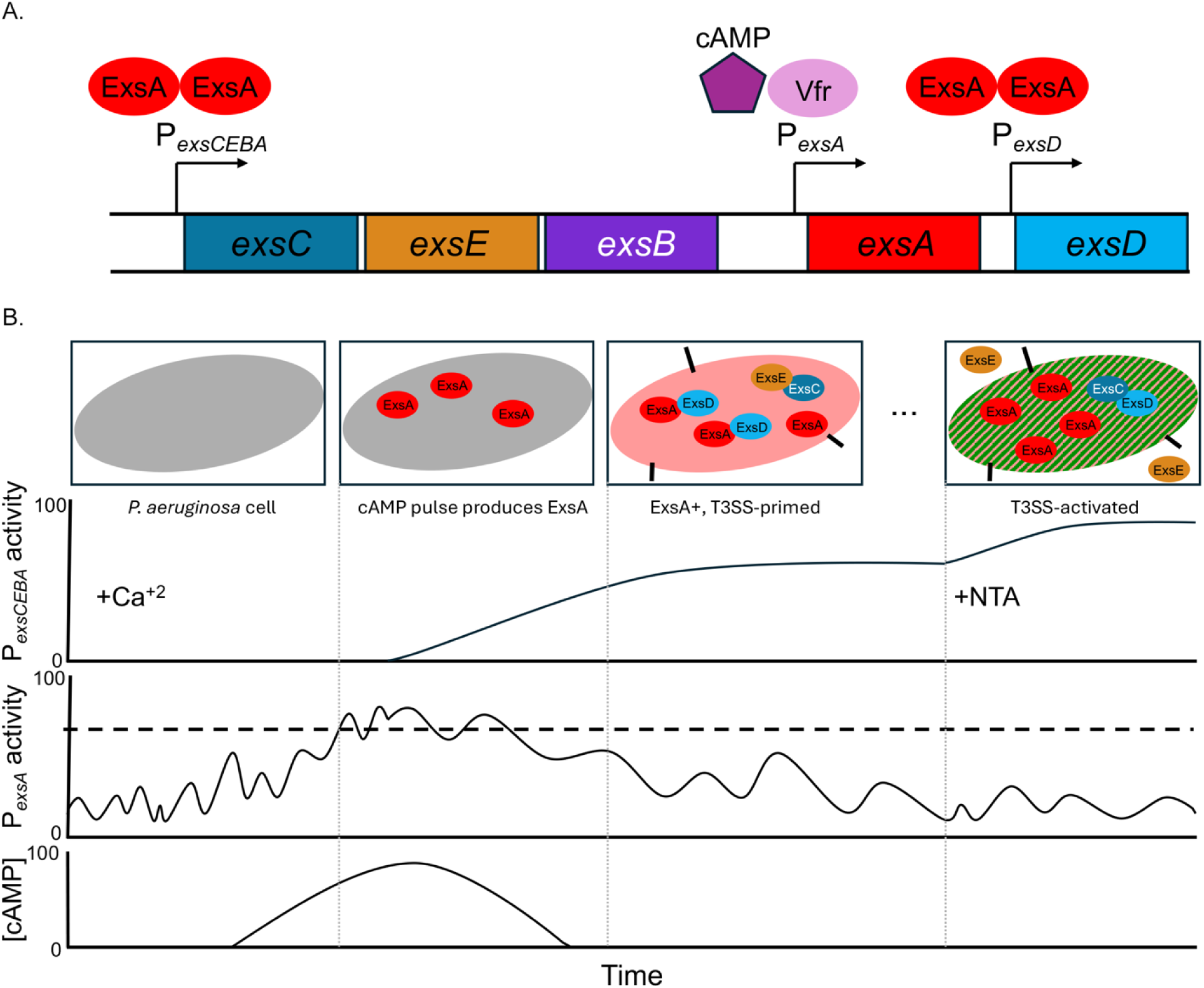
The cAMP-responsive P*_exsA_* promoter coupled with positive feedback at P*_exsCEBA_* creates a bistable distribution of primed cells. (A) T3SS regulatory gene schematic including ExsA-dependent and ExsA-independent promoters. (B) Proposed model for generating a bistable ExsA+ cell with assembled T3SS injectisome via cAMP-responsive P*_exsA_* expression. A *P. aeruginosa* cell with below threshold P*_exsA_* activity responds to a pulse of increased cAMP by producing ExsA from the P*_exsA_*promoter. Increased ExsA leads to a burst of ExsA-dependent gene expression, resulting in a “primed” *P. aeruginosa* cell with an assembled, “closed” T3SS injectisome; ExsA is bound to and inhibited by ExsD. In response to a T3SS-activating signal, the assembled T3SS opens and ExsE is secreted, leaving ExsA free to dimerize and committing the cell to T3SS gene expression.

cAMP is a well-recognized positive regulator of T3SS gene expression (25, 36, 38) whose levels are determined by activity of the adenylate cyclases CyaA and CyaB and the phosphodiesterase, CpdA (36, 48). We used a well-characterized transcriptional reporter, *lacP1*-sfGFP, to examine [cAMP] at the single cell level across different *P. aeruginosa* strains that vary in the proportion of cells that are T3SS-ON. Strains that had higher GFP MFI – PA103 and PAK – also had high proportions of cells that were T3SS-ON, while the lower [cAMP] levels observed in PA14 and PAO1 cells were correlated with bimodal distributions of T3SS-OFF and -ON cells. These findings are consistent with our model, in which higher [cAMP] increases the likelihood that a cell will become primed. PA103 cells were also remarkable for the high proportion of bacteria with assembled needles, while <20% of PA14 cells had needles by cryo-ET. Although many mechanism(s) could generate interstrain variation in intracellular [cAMP], overexpression of the CpdA phosphodiesterase from a plasmid was sufficient to decrease the population of primed cells for both PA14 and PA103.

cAMP production is mechanistically linked to T4P assembly/retraction via the Pil/Chp chemotaxis system, which activates the CyaB adenylate cyclase (38). Our experiments were carried out with bacteria that were cultured planktonically; under these conditions, many PA103 bacteria assembled T4P (as imaged by whole cell cryo-ET) while this was rarely observed for PA14 cells. We introduced the P*_exoT_* reporter into a series of PA103 and PA14 mutants disrupted for T4P assembly or retraction, and observed little effect on the proportion of T3SS-ON cells, with a few notable exceptions. Specifically, disruption or deletion of the PilJ chemoreceptor significantly reduced the population of T3SS-ON cells, as did mutation of PilA, which is proposed to interact with and activate PilJ (39).

PilJ activation leads to increased production of cAMP by the CyaB adenylate cyclase; this interaction requires FimL and is modulated by the methylation state of the PilJ (55). The deletion of *pilJ*, *fimL* or *pilA* significantly decreased priming in the PA14 dual reporter strain. Conversely, an in-frame deletion of the PilJ periplasmic “ligand binding domain” (PilJΔLBD1-2) markedly increased the proportion of primed cells. Further deletion of *pilA* in this background (PilJΔLBD1-2 Δ*pilA*) diminished priming but not to the level seen for Δ*pilA*, arguing that this mutant PilJ allele has increased basal activity that does not require interactions with PilA. Deletion of the methylesterase *chpB* also increased priming, consistent with increased adenylate cyclase activation by methylated PilJ (55). Again, deletion of *pilA* in this background (Δ*chpB* Δ*pilA*) reduced the population of primed cells, but to a level that was still elevated relative to WT or Δ*pilA*.

Surface-attached *P. aeruginosa* increase [cAMP] in a T4P-dependent manner(39, 59). Coupling T3SS expression to surface attachment could provide *P. aeruginosa* with a defense mechanism against eukaryotic predators when it is at a surface but not yet within a biofilm community (38, 60). We grew our dual reporter strain and isogenic mutants on agar surfaces and observed that the deletion of *pilA*, *pilJ* or *fimL* reduced primed cell populations to a far greater extent than we observed for liquid grown cells (Fig 6G). The PilJΔLBD allele strongly increased the proportion of primed cells under these conditions, and – as we observed for liquid-grown cells – deletion of *pilA* in this background diminished the proportion of primed cells but did not return it to wild-type or Δ*pilA* levels. Because we could not rigorously control the amount of Ca^+2^ present in agar plates, we did not compare the proportions of primed cells between surface vs. liquid grown cells in our experiments.

Though T3SS regulation differs among bacterial pathogens, many species exhibit bimodal T3SS expression (7, 61). In *Salmonella* Typhimurium, T3SS-1 expression is controlled by a complex ‘feed-forward’ loop of multiple positive autoregulatory elements. The strongest of these, HilD, directly promotes the expression of HilA, the T3SS-1 transcriptional activator, while two other positive autoregulators (HilC and RtsA) serve as rheostatic amplifiers for HilA (62). HilD is repressed by binding to HilE, the main negative T3SS-1 regulator; this interaction primarily controls the bistable switch and sets the threshold for generating a T3SS-1 ‘ON’ cell (62, 63). Unlike *P. aeruginosa’s* ExsD, HilE expression is not controlled by HilA, but rather by PhoPQ (64). *S.* Typhimurium T3SS-1 expression is bistable, though hypoxia – which inactivates HilE - can cause a shift to unimodal T3SS-1+ expression (65). As additional transcriptional activators for the T3SS-1 are downstream of HilA, the *S.* Typhimurium system acts in a multi-step, hierarchical process with the potential to create ‘primed cells’ (62). Small increases in HilD, combined with inputs from HilC and RtsA, could shift the balance between HilA and HilE, resulting in expression of T3SS structural genes before cells receive the T3SS-activating signal of host-cell contact (62). Heterogeneous T3SS expression is not limited to mammalian pathogens, but is also found in plant pathogens including *Dickeya dadantii* and *P. syringae*, suggesting that T3SS-priming could be more widespread if it is a key feature of bistability (66, 67).

To our knowledge, no single cell studies of T3SS expression have been carried out in Vibrio *spp.* to determine if T3SS expression is bimodal. *V. alginolyticus,* an opportunistic marine animal and human pathogen, does utilize a partner-switching mechanism to control T3SS gene expression and requires ExsE secretion for T3SS activation and upregulation (68). This requirement for ExsE secretion could suggest that T3SS needles are assembled prior to receiving a secretion signal, similar to our observations in *P. aeruginosa*. The related species, *V. parahaemolyticus* also regulates its T3SS1 through partner-switching, but is not reliant on ExsE secretion to increase T3SS-mediated virulence (33, 68-72). Crucially, ExsA does not regulate its own expression in either of these species, eliminating one known mechanism for establishing bistability (73).

In conclusion, the T3SS is a critical virulence factor for *P. aeruginosa* whose expression carries significant fitness costs for individual bacteria. The ability to express the transcriptional activator, ExsA, independently of ExsD, ExsC and ExsE from the P*_exsA_* promoter is critical for T3SS expression: deletion of Vfr markedly reduces the proportion of bacteria expressing T3SS gene just as we report for bacteria lacking both adenylate cyclases (Figure 4), while disruption of the P*_exsA_* promoter itself completely ablates T3SS expression (25, 74). These findings likely reflect the importance of ‘noisy’ transcription to the generation of primed cells, from a promoter that can also respond to the critical second messenger cAMP.

## Materials and Methods

### Bacterial strains and growth conditions

The bacterial strains and plasmids used in this study are listed in Table S1. Cultures were routinely grown at 37°C with shaking in Miller’s Luria Broth (LB) consisting of 10 g/L casein digest peptone, 10 g/L sodium chloride, and 5 g/L yeast extract unless otherwise indicated. For biparental matings, *P. aeruginosa* was selected on Vogel-Bonner Medium (VBM) agar (75). When necessary, antibiotics were added at the following concentrations: for *E. coli*, 15 μg/mL gentamicin, 20 μg/mL tetracycline, 100 µg/mL ampicillin; for *P. aeruginosa*, 30 µg/mL (CRISPR mating, *attTn7* complementation) or 100 µg/mL gentamicin, 100 µg/mL tetracycline, 200 µg/mL carbenicillin.

To activate T3SS gene expression, *P. aeruginosa* was grown in MinS medium with 10 mM nitrilotriacetic acid (NTA) added to MinS in selected experiments (26). To repress T3SS gene expression, *P. aeruginosa* was grown in MinS medium plus 5 mM CaCl_2_ (MinS + Ca^+2^) (76). As indicated, cyclic-AMP (cAMP) (MedChemExpress) dissolved in 50 mM Tris-HCl (pH = 8) was added to liquid medium at a final concentration of 20 mM.

### Plasmid and strain construction

Reporters for ExsA (*exsA*-IRES-mTagRFP-t) and T3SS gene expression (*attB*::P*_exoT_*-sfGFP) were previously constructed and introduced to *P. aeruginosa* as detailed (26). To report relative intracellular levels of cAMP, a *lacP1*-sfGFP reporter was introduced to *P. aeruginosa* at the *attB* site through biparental mating (38, 77). Mating was carried out on low-salt LB agar plates (containing 5 g/liter NaCl), and integrants were selected on VBM-tetracycline plates. The vector backbone was excised by mating with *E. coli* SM10 pFLP2, which expresses Flp recombinase (78). Successful loss of vector backbone and pFLP2 was confirmed by PCR for candidates that were both tetracycline and carbenicillin sensitive. The *pilJ* gene was deleted through biparental mating of the previously generated PA14 dual reporter with a Gateway adapted pDONRx plasmid containing ∼1kb up- and down-stream flanks for *pilJ* as previously described, with merodiploid integrants selected on VBM-gentamicin plates and counterselection on VBM plates containing 15% (w/v) sucrose to promote backbone loss (38). Δ*pilA*, *ΔchpB, ΔchpB ΔpilA, ΔpilK*, *ΔfimL*, *pilJ* ΔLBD1-2, and *pilJ* ΔLBD1-2 Δ*pilA* constructs were generated using the CRISPR-Cas9 system as previously described (79). PAM sites and single guide RNA sequences were identified using Geneious Prime 2025.2. (http://www.geneious.com). pS648 plasmids with the target sequence and pSH124-*ssr* were independently introduced to the PA14 dual reporter through electroporation, and plated on LB plates containing carbenicillin, gentamicin, and 5 mM m-toluic acid. The *pilJ* deletion mutant was complemented chromosomally at the *att-*Tn7 insertion site as previously described through biparental mating of PA14 Δ*pilJ attB::*P*_exoT_*-*sfGFP exsA-*IRES-*mTagRFP-t* with pUC18T-*pilJ*, and helper conjugation plasmids pTSN2 and pRK2013 (80, 81). The pUC18T-*pilJ* construct was generated through PCR amplification of WT PA14 *pilJ* sequence, including the native ribosome binding site as well as homologous overhangs to pUC18T, then assembled using HiFi Assembly (NEB) on a SacI/HindIII digested pUC18T. All constructs were validated through Sangar sequencing (Keck Sequencing Facility, Yale). Primers for this study, including spacer guide sequences, are listed in Table S2.

### Flow cytometry

Single bacterial colonies were inoculated into LB and grown overnight with aeration at 37°C, subcultured into LB and grown until early log-phase, washed with MinS, and diluted to a starting OD_600_ of 0.01 into the indicated MinS medium. For surface-grown bacteria, bacteria were spread at an OD_600_ of 0.01, on 25 mL MinS agar plates. Cultures were sampled at 7 hours postinduction and processed for flow cytometry. For surface experiments, the entire agar plate was scraped and resuspended into PBS-MC (Phosphate Buffered Saline pH 7.4 plus 0.9 mM calcium, 0.9 M magnesium), then processed for flow cytometry as previously detailed (82). Briefly, bacterial cells were fixed with 1% paraformaldehyde with aeration at 37°C for 30 minutes and diluted to an OD_600_ of 0.04 in PBS-MC, then passed through a 40 µm filter. Cells were analyzed for GFP emission in B525-FITC with a 525/40 bandpass filter and RFP emission in 584-mCherry with a 584/42 bandpass filter using an CytoFlex LX equipped with a 96-well plate sampler (Beckman Coulter). At least 50,000 events were collected per sample. Isogenic WT *P. aeruginosa* bacteria lacking fluorescent reporters were used as gating controls as they showed equal fluorescence to T3SS-null PA14 Δ*exsA* P*_exoT_*-sfGFP (Figure S11). The full gating scheme is detailed in Figure S12.

### Cryo-electron tomography sample preparation and data analyses

*P. aeruginosa* strains PA103 and PA14 were grown as described above, either in MinS + 5 mM Ca^+2^ or MinS + 10 mM NTA media as indicated, with either 20 mM cAMP or Dulbecco-PBS (D-PBS) added. After growth, bacterial concentrations were adjusted to OD_600_ 1.0 in DPBS. Bacterial samples and BSA-coated gold tracer (10nm) solution (Aurion) were then mixed at a ratio of 1:1 (V/V). 5μL of the mixture was deposited onto freshly glow-discharged cryo-EM grids (Quantifoil, Cu, R2/1, 200 mesh). Filter paper (Whatman^TM^) was used to blot the sample for ∼5 seconds from the back of the cryo-EM grid, which was then immediately plunged into liquid ethane and propane mixture using a homemade gravity plunger. Frozen-hydrated specimens of *P. aeruginosa* were imaged at ∼-180°C using Titan Krios G2 300kV transmission electron microscope (Thermo Fisher Scientific) equipped with a field emission gun and a K3 direct detection camera with a GIF BioQuantum Imaging Filter (Gatan). SerialEM software was used to record tilt series images for each bacterial target at a magnification of 15,000 (83). The physical pixel size at the specimen level is 6.184Å. The angle of the tilt series ranged between ±48° in 3° increments. The total electron dose was ∼60e^-^/Å^2^ distributed across 33 images of each tilt series. The defocus was set at - 10 μm without volta phase plate and -1.0 μm with phase plate (Table S3). MotionCor2 was used to correct image drift during data collection (84). Drift-corrected images were combined to create image stacks for each tilt series by IMOD (85). Fiducial beads in the images were used to align tilt series by IMOD (85). The binvol function in IMOD was used to generate 4x binning of the aligned stacks, and 4x binned tomograms with SIRT reconstruction were then reconstructed using Tomo3D (86). The number of tomograms for each specimen is shown in Table S3. Injectisomes on the bacterial surface were counted manually, and the graph was created using GraphPad Prism, version 10.

### Statistical Analysis

GraphPad Prism software version 10.5.0 was used for all statistical analysis. Two-way comparisons were performed using two-tailed unpaired *t* tests with Welch’s correction to compare means for normally distributed data sets without assuming populations have equal variance. Multiple comparisons for normally distributed data were carried out using one-way ANOVA with Dunnett’s multiple comparisons test.

